# Pharmacokinetics and Pharmacodynamics of Vancomycin derivative LYSC98 in a Murine Thigh Infection Model against *Staphylococcus aureus*

**DOI:** 10.1101/2021.12.14.472732

**Authors:** Peng He, Xin Li, Xiaohan Guo, Xingchen Bian, Meiqing Feng

## Abstract

**LYSC98** is a vancomycin derivative used for gram-positive bacterial infections therapy. We reported the pharmacokinetic/pharmacodynamic (PK/PD) targets of LYSC98 against *Staphylococcus aureus* using a murine thigh infection model. Three *Staphylococcus aureus* strains were utilized. Single-dose plasma pharmacokinetics of LYSC98 were determined in infected mice after the tail vein injection of 2, 4, and 8mg/kg. The results showed maximum plasma concentration (C_max_) 11466.67 - 48866.67 ng/mL, area under the concentration-time curve from 0 to 24 h(AUC_0-24_) 14788.42 -91885.93 ng/mL·h, and elimination half-life(T_1/2_) 1.70-2.64 h, respectively. The C_max_ (R^2^ 0.9994) and AUC_0-24_ (R^2^ 0.981) were positively correlated with the dose of LYSC98 in the range of 2-8 mg/kg.

Dose fractionation studies using total doses of 2 to 8 mg/kg administered with q6h, q8h, q12h, and q24h were performed to evaluate the correlation of different PK/PD indices with efficacy. Sigmoid model analysis showed C_max_/MIC (R^2^ 0.8941) was the best PK/PD index to predict the efficacy of LYSC98. In the dose ranging studies, two Methicillin-resistant *Staphylococcus aureus* (MRSA) clinical strains were used to infect the mice and 2-fold-increasing doses (1 to 16 mg/kg) of LYSC98 were administered. The magnitude of LYSC98 C_max_/MIC associated with net stasis, 1, 2, 3 and 4 - log10 kill were 5.78, 8.17, 11.14, 15.85 and 30.58, respectively. The results of this study showed LYSC98 a promising antibiotic with in vivo potency against MRSA, and will help in the dose design of phase one study for LYSC98.

## INTRODUCTION

*Staphylococcus aureus* is a Gram-positive human commensal which inhabits approximately 30% of healthy people in the anterior nares (Lowy, 1998; Rao et al., 2015). As a leading cause of hospital-associated (HA) and community-associated (CA) bacterial infections, *S. aureus* is associated with numerous mild skin and soft tissue infections, as well as life-threatening pneumonia, bacteremia, osteomyelitis, endocarditis, sepsis and toxic shock syndrome(David and Daum, 2010; Zhou et al., 2018).

Penicillin remains the drug of choice if the isolate is sensitive to it(Drebes et al., 2014). But once bacterial resistance occurs, such as methicillin-resistant *Staphylococcus aureus* (MRSA) infections, vancomycin is the advanced drug to treat(Lakhundi and Zhang, 2018). Patients who cannot tolerate vancomycin have been treated with fluoroquinolones, trimethoprim - sulfamethoxazole, clindamycin, or minocycline. Each of these drugs is effective in cases where sterilization is required. However, they are not as effective as vancomycin, either because they have a less anti-staphylococcal activity or because drug resistance develops during treatment(Chambers, 1997; Lakhundi and Zhang, 2018; Michel and Gutmann, 1997). Alternatively, *S. aureus* clinical isolates with reduced susceptibility to vancomycin, and less commonly, with complete resistance to vancomycin have emerged within the past 20 years(Hidayat et al., 2006; Howe et al., 1998; McGuinness et al., 2017). Vancomycin-resistant *S. aureus* (VRSA) are associated with persistent infections, vancomycin treatment failure, and poor clinical outcomes(McGuinness et al., 2017).

LYSC98 is a new synthesized compound derived from vancomycin by chemical modification of its side chain (**Fig.1**). In our previous study, it has been confirmed that LYSC98 has a similar antibacterial spectrum to vancomycin and other glycopeptide antibiotics, but is higher than vancomycin in the intensity and durability of antibacterial activity, especially the activity against vancomycin-resistant *S. aureus* (Supplementary Table 1). In addition, LYSC98 showed excellent in vivo protective effect against *S. aureus* infection in mice. Its ED50 is 0.41-1.86 mg/kg, which is better than vancomycin (2.32-5.84 mg/kg) and linezolid (3.07-7.60 mg/kg) (Supplementary Table 2). However, there were no studies designed to identify the PK/PD index and target values related to the efficacy of LYSC98. PK/PD studies become an indispensable part of antibacterial drug development. PK/PD targets are used to support the dosing regimen design in phase I clinical trials and are necessary data for the susceptible breakpoints establishment(Mouton et al., 2012). Animal PK/PD studies are frequently employed in the determination of PK/PD targets because of the consistent target results to that in humans, as well as the flexibility dosing regimen design to analyze exposure-response relationships.

**FIGURE 1.**
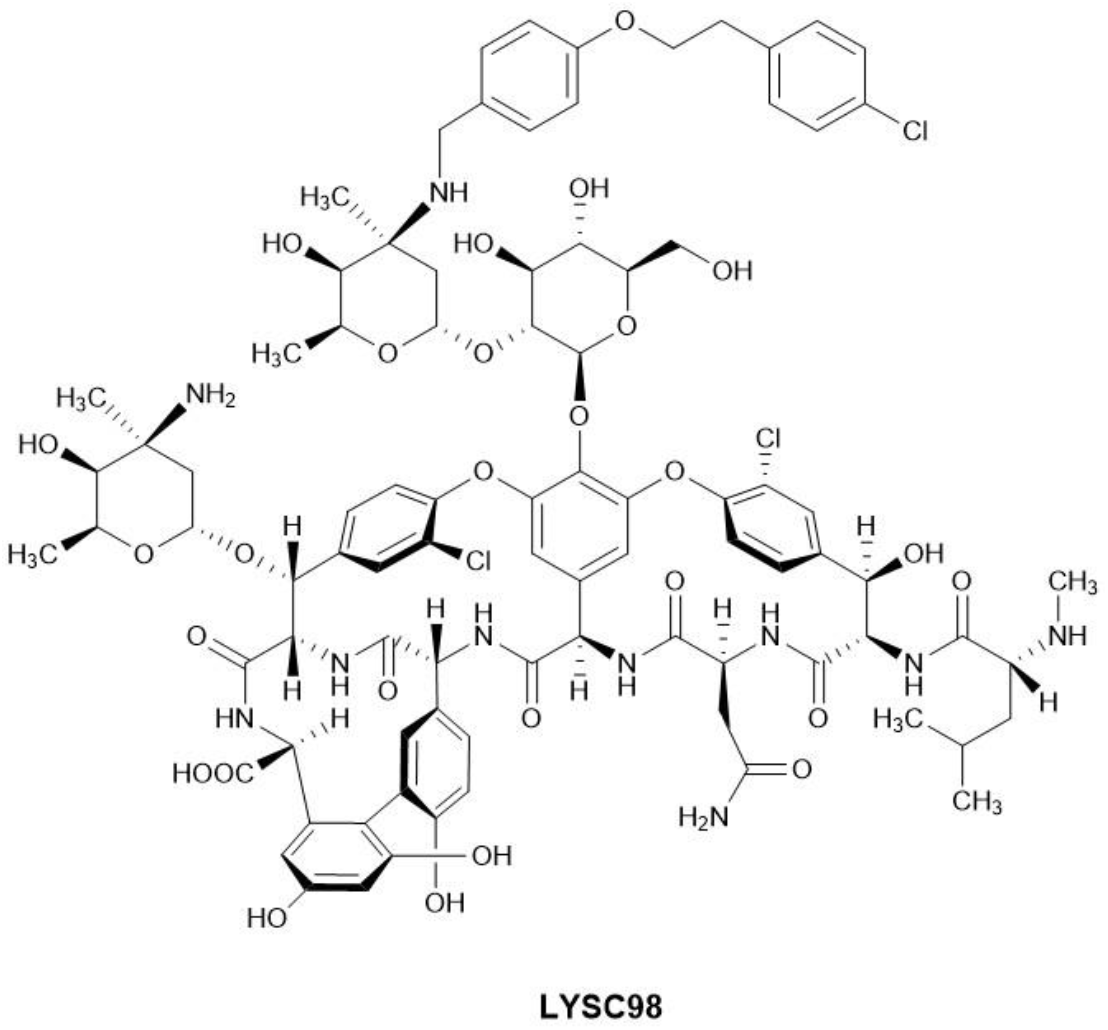
The chemical structure of LYSC98.

In this study, we used an immunosuppressed murine thigh infection model against various *S. aureus* strains to identify the pharmacokinetic and pharmacodynamic characteristics of LYSC98. The studies were designed to determine the PK profile in the murine infection model, the exposure-response relationships and PK/PD targets of LYSC98 associated with efficacy against *S. aureus.* We hope to provide a basis for the formulation of clinical administration plan and drug sensitive breakpoint for LYSC98.

## MATERIALS AND METHODS

### Bacterial, Media, and Antibiotic

Two clinical MRSA strains 18-W26-14 and 18-W27-73[Institute of Antibiotics, Huashan Hospital, Shanghai] and one reference *S. aureus* strain ATCC29213 were used in this study (**Table 1**). Bacteria were cultured and quantified on a LB agar plate and grown for 16 h at 37°C. LB agar plate consists of 1% Peptone (OXOID), 0.5% yeast powder (OXOID), 1% sodium chloride (Sinopharm), and 1.5 AGAR powder (Meilunbio, Dalian). LYSC98 (Purity:92.02, Lot no:20200322) was supplied by Shanghai Laiyi Center for Biopharmaceutical R&D (Shanghai, China). The compound was reconstituted and diluted to appropriate concentrations with 5% glucose solution.

**TABLE 1.** In vitro activity of LYSC98 against study organisms.

| Organisms | Description | LYSC98 MIC (mg/L) |
| --- | --- | --- |
| ATCC29213 | - | 2 |
| 18-W26-14 | MRSA | 4 |
| 18-W27-73 | MRSA | 2 |

MRSA, methicillin-resistant *Staphylococcus aureus*.

### *In Vitro* Susceptibility Testing

The minimum inhibitory concentration (MIC) of LYSC98 against all isolates were determined in triplicate with broth microdilution method according to Clinical and Laboratory Institute (CLSI, 2018) guidelines. ATCC 49619 was served as a quality control strain. ATCC 29213 was served as a quality control strain.

### Murine Thigh Infection Model

The animal studies were approved by the Experimental Animal Ethics Committee of Pharmacy in Fudan University and followed the Experimental Animal Welfare Review Guide. Six-week-old, specific-pathogen-free, male ICR mice (SLAC Laboratory Animal Co., Ltd, Shanghai) weighing 18–22 g was used in all studies. The neutropenic murine thigh infection model was established as previously described(Growcott et al., 2019). Animals were rendered neutropenic by intraperitoneal injections of 150 and 100 mg/kg cyclophosphamide (Sigma–Aldrich, St Louis, MO, USA), on days −4 and - 1 respectively prior to infection. Two hours prior to treatment (−2 h), 200μL containing about 1-2* 10^6^ CFU of bacterial inoculum was administered into the bilateral gastrocnemius muscle via an intramuscular injection. At 0 h a cohort of animals was sacrificed via CO2 to determine the bacterial levels at the start of treatment. The remaining animals were euthanized 24 h after the start of therapy. The infected thighs were excised and homogenized in sterile saline (0.9% w/v) until the tissue was completely homogenized. The homogenates were serially diluted in sterile saline before dilutions were plated on LB Agar plates, incubated overnight at 37°C and the colonies counted. The CFU/thigh were calculated and transformed to log_10_.

### Pharmacokinetics

Single-dose pharmacokinetics of LYSC98 was studied in thigh infected mice at doses level of 2,4 and 8 mg/kg following intravenous administration (0.2 ml/dose). Blood samples were collected at 0.25, 0.5, 1, 2, 4, 8, 12, 18, 24 h at each dose. Three mice were used per time point. Plasma was separated by centrifugation at 4000 g for 10 min at 4°C and stored at −80°C until urea and LYSC98 concentration analysis. The detection of compound concentration was entrusted to Shanghai Institute of Materia Medica, Chinese Academy of Sciences.

WinNonlin software (Version 6.3; Pharsight Inc., St. Louis, MO, United States) was employed to calculate the PK parameters using a noncompartmental model, including the elimination half-life (t_1/2_), the area under the concentration-time curve over 24 h (AUC_0-24_), and the peak drug concentration (C_max_). The PK parameter estimation for treatment doses that were not directly determined was calculated based on a compartment model.

### Pharmacokinetic/Pharmacodynamic Index Determination

Neutropenic mice were infected with the standard strain of *S. aureus* ATCC29213 for a dose fractionation experiment. Treatment with LYSC98 by the intravenous route was initiated 2 h after inoculation. Dose-fractionation study is useful in reducing the interdependence among the PK/PD index and confirming which one is the most important for efficacy. The total daily doses included 2, 4, 8mg/kg, divided evenly every 6, 8, 12 and 24 h. Groups of three mice and six thighs were included in each dosing regimen. The mice were sacrificed 24 h after the first dosing of LYSC98, bilateral thigh muscles were aseptically removed and processed for CFU determination(Asempa et al., 2020; Lepak et al., 2019). Briefly, the thigh muscles were homogenized in 10-fold volumes of 0.9% NaCl under 4 °C. Serial 1:10 dilutions of the homogenate were plated overnight at 37°C for CFU counting. The lower limit of viable colony counts was 2000 CFU/ml. Results were expressed as the mean number of log_10_CFU per gram tissue for six samples. Untreated mice used for the growth control assessment were similarly sacrificed before treatment and 24 h after treatment. Efficacy was calculated as the change in the log_10_CFU obtained at 24 h for LYSC98-treated mice.

To determine the dominant PK/PD index driving efficacy, the number of bacteria in the thigh muscles at the end of therapy was correlated with three parameters: the free drug peak level divided by the MIC (*f*C_max_/MIC), the area under the free concentration-time curve over 24 h divided by the MIC (*f*AUC_0-24_/MIC), and the cumulative percentage of a 24 h period that the free drug concentration in plasma exceeds the MIC (%*f*T > MIC), for each of the dosage regimens studied. The mathematical model used was derived from the Sigmoid E_max_ model:

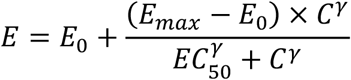

where E is the effect, in this case, the log_10_ change in CFU per gram thigh muscles between treated mice and untreated controls after the 24-h period of study, E_max_ is the maximum effect, C is the PK/PD index value, EC50 is the value of PK/PD index required to achieve 50% of the E_max_, and γ is the slope of the dose-effect curve. The R^2^ value from non-linear regression analysis (WinNonlin 6.3) was used to assess the correlation of treatment efficacy with each of the PK/PD indices.

### PK/PD Targets for Efficacy

Dose-ranging efficacy studies were then performed to determine the PK/PD targets for net stasis, 1-log_10_CFU, 2-log_10_CFU, 3-log_10_CFU and 4-log_10_CFU kill with two clinical *S. aureus* strains using the murine neutropenic thigh infection model. Increasing single dosing regimens of LYSC98 were administered varied from 1 to 16 mg/kg. Groups of three mice were included per dose level. Treatment was initiated 2 h after inoculation. Animals were sacrificed at 24 h after therapy, and the bilateral thigh muscles were aseptically removed and immediately processed for CFU determination. A sigmoid dose-response model derived from the Hill equation was used to calculate the dose, and PK/PD targets of LYSC98 producing a net bacteriostatic effect, 1- log_10_CFU, 2-log_10_CFU, 3-log_10_CFU and 4-log_10_CFU kill over 24 h compared to the organism burden at the start of treatment.

## RESULTS

### *In Vitro* Susceptibility Testing

The MIC values of LYSC98 against three *S. aureus* isolates used in this study are shown in **Table 1**, ranging from 2 to 4 mg/L.

### Pharmacokinetics

Single-dose PK of LYSC98 in plasma are shown in **Figure 2**. The elimination halflife in plasma ranged from 0.25 to 0.33 h. C_max_ concentrations ranged from 11466.67 to 48866.67 ng/mL and were linear across 2-8 mg/kg dose range (R^2^ 0.9994). AUC_0-24_ values ranged from 14788.42 to 91885.93 ng/mL and were linear across 2-8 mg/kg dose range (R^2^ 0.981). Detailed PK parameters are listed in **Table 2.**

**FIGURE 2.**
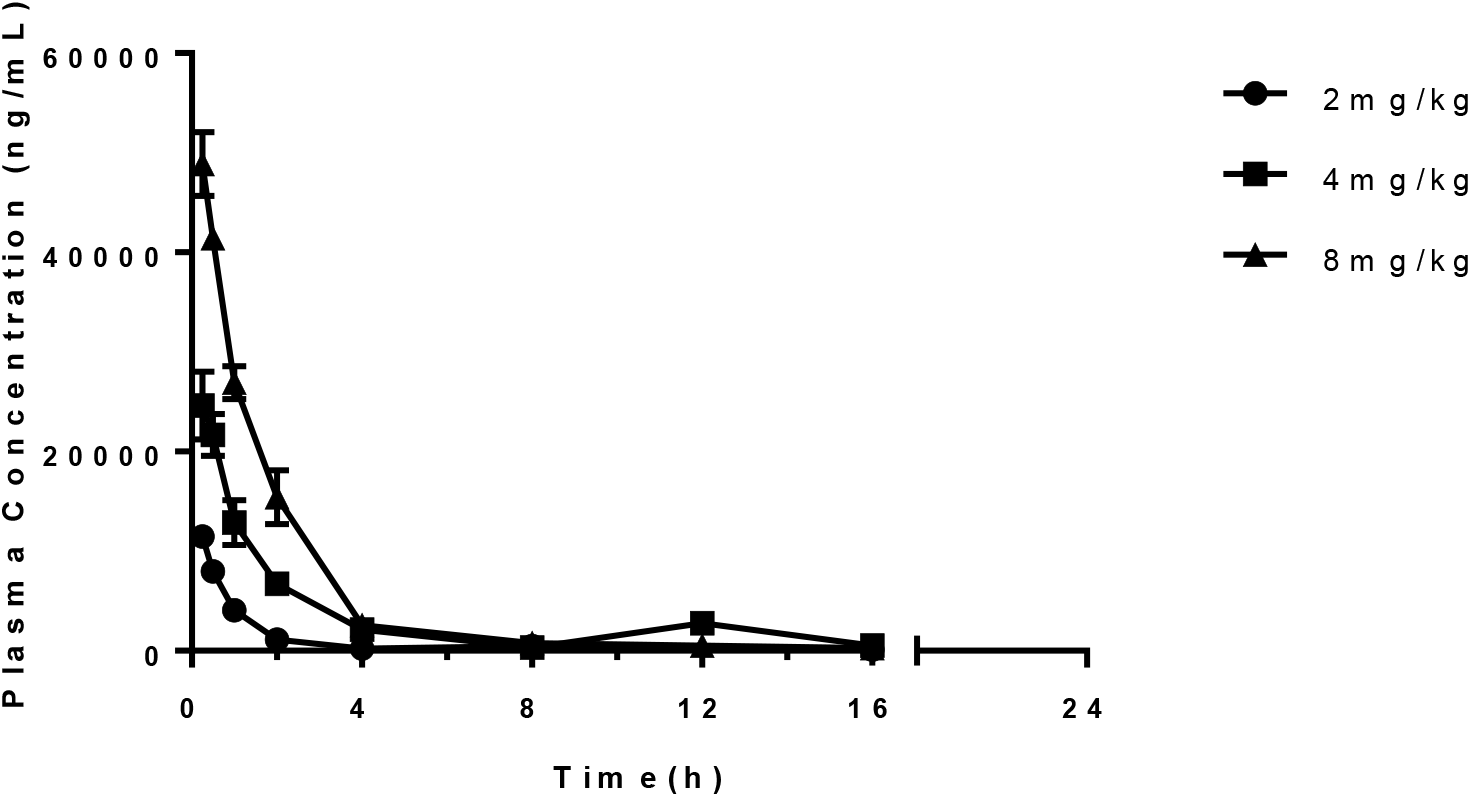
Pharmacokinetics of LYSC98 in plasma following single intravenous doses at 2-8 mg/kg in neutropenic thigh infected mice. Groups of three mice were sampled for each time point. Each symbol represents the mean value of three mice. The error bar represents the standard deviations.

**TABLE 2.**
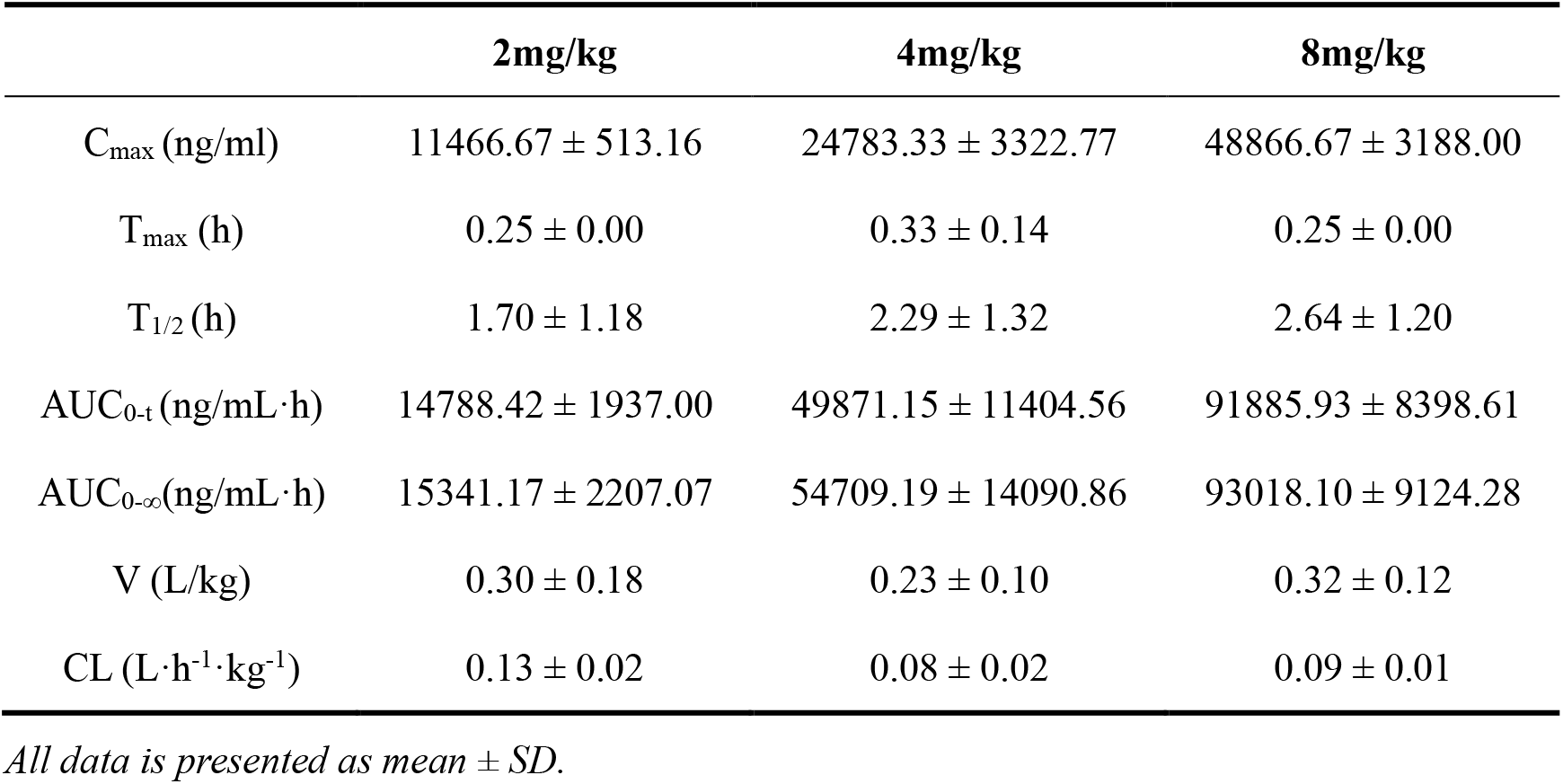
PK parameters calculated using non-compartment model

### PK/PD Index Determination

LYSC98 treatment produced mostly 5.02 log_10_ reduction of CFU burden against *S. aureus* ATCC29213 in the dose fractionation experiment (**Fig3**). The dose-response curves with different dosing intervals showed the bactericidal effect was improved with the increase of dose but not the decrease of dosing interval. The relationship between efficacy and three PK/PD indices *f*AUC_0-24_/MIC, %*f*T > MIC and *f*C_max_/MIC are shown in **Figure 4**, and the value of the square of the correlation coefficient (R^2^) were 0.7793, 0.5424 and 0.8941, respectively (**Table 3**). We considered *f*C_max_/MIC as the PKPD index of LYSC98 since its best correlation with efficacy.

**FIGURE 3.**
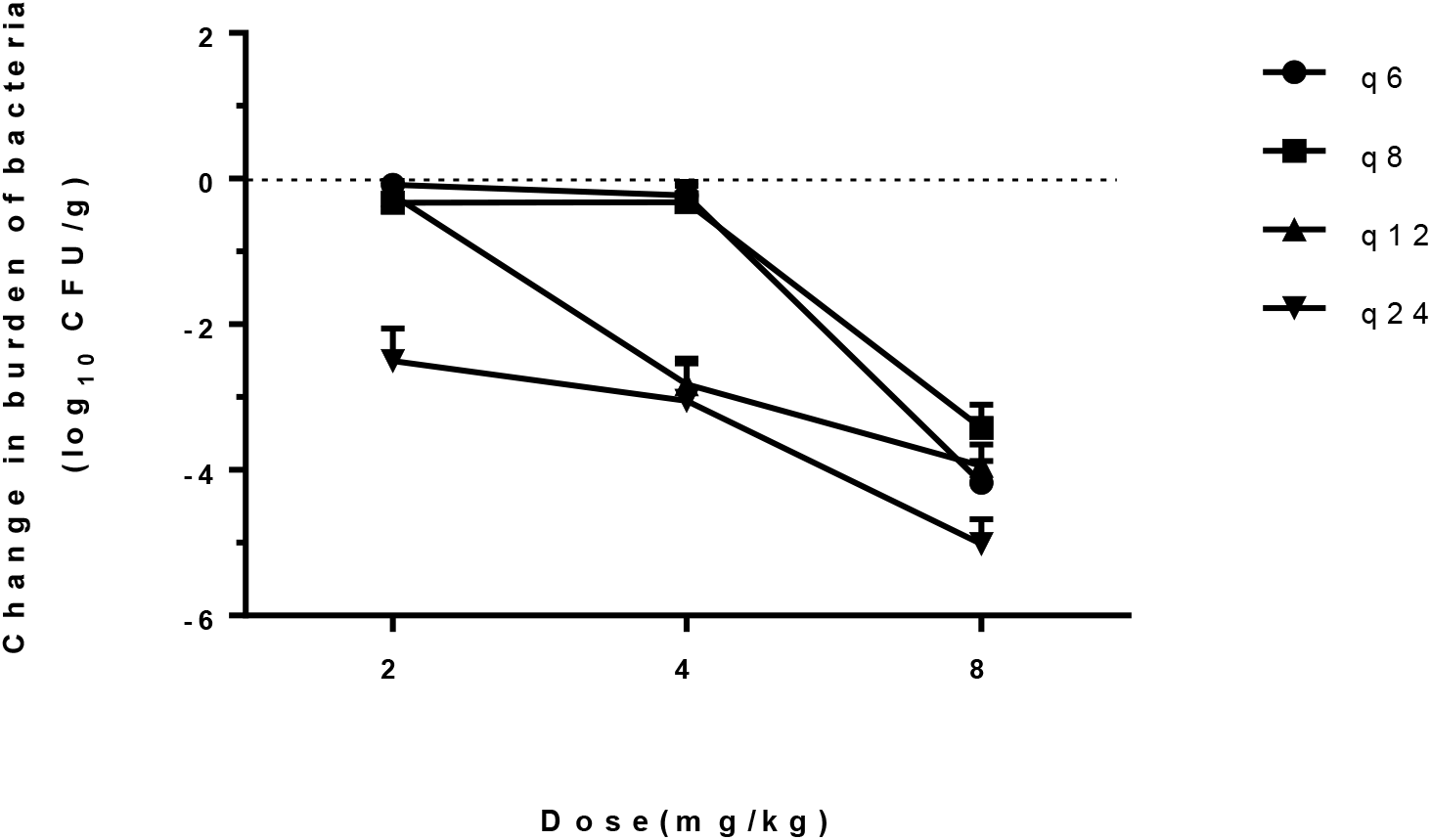
Regimens of LYSC98 treatment produced CFU burden reduction against *S. aureus* ATCC29213 in the dose fractionation experiment. Abscissa q6, q8, q12, and q24 means LYSC98 were treated per 6,8,12 and 24 hours under constant total dose in each group. Each symbol represents the mean value of three mice. The error bar represents the standard deviations.

**FIGURE 4.**
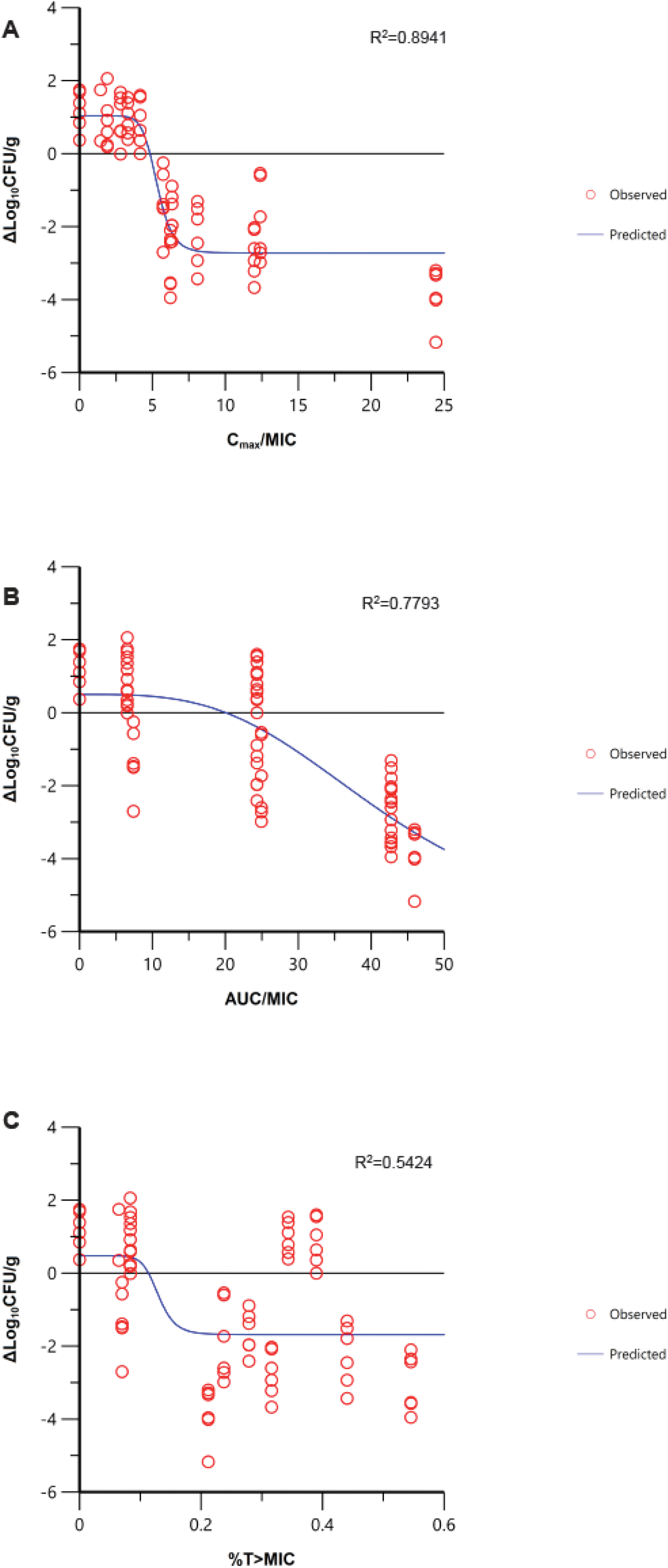
Correlation of pharmacokinetic/pharmacodynamic (PK/PD) indices *f*AUC_0-24_/MIC (A), %*f*T > MIC (B) and *f*C_max_/MIC (C) with efficacy of LYSC98 in a neutropenic murine thigh infection model caused by *S. aureus* ATCC29213 in dose-fractionation study. Treatment was initiated at 2 h post infection. LYSC98 was intravenously administered with a dosing range of 2-8 mg/kg, in once daily (q24h), twice daily (q12h), three times a day (q8h) and four times a day (q6h). Each circle represents data for each mouse.

**TABEL 3.** E_max_ model parameters characterizing the relationship between efficacy and different PK/PD indices of LYSC98

| Parameters | $E_{max}$ | $E_0$ | $EC_{50}$ | $\gamma$ | $R^2$ |
| --- | --- | --- | --- | --- | --- |
| $C_{max}/MIC$ | -3.76 | 1.04 | 5.32 | 9.99 | 0.8941 |
| $AUC_{0-24h}/MIC$ | -6.79 | 0.50 | 42.90 | 3.34 | 0.7793 |
| %T>MIC | -2.16 | 0.47 | 0.13 | 9.39 | 0.5424 |

### PK/PD Targets for Efficacy

Two additional *S. aureus* strains were used in the dose-escalation experiment to determine the PK/PD targets required for efficacy.. The dose-response data for the two strains are shown in **Figure 5**.

**FIGURE 5.**
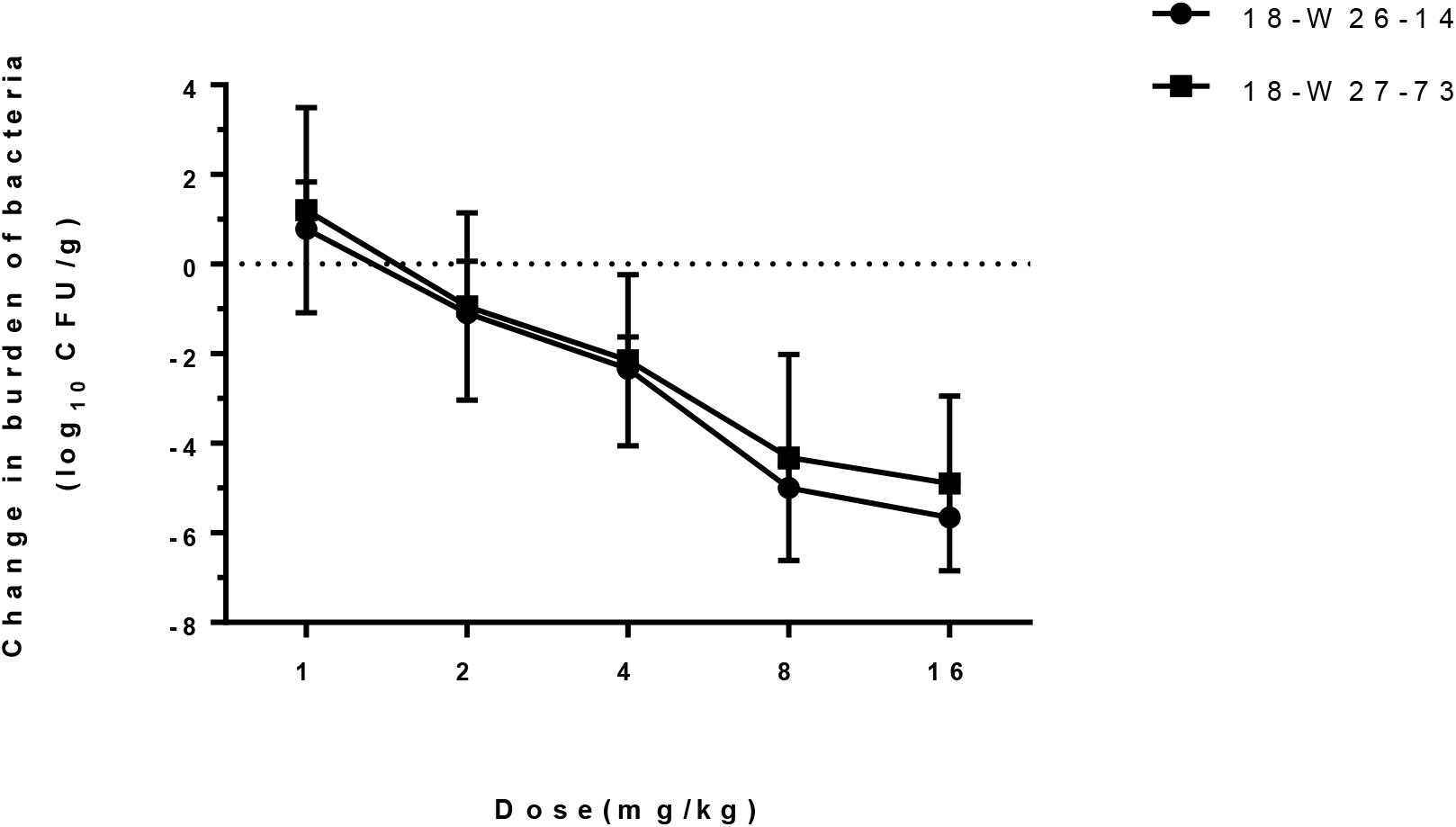
In vivo dose-response curves for LYSC98 against two *S. aureus* strains over a 24 h study period after a single dose administration in the neutropenic murine thigh infection model. Each symbol represents the mean and standard deviation from three mice. The burden of organisms was measured at the start and end of therapy. The horizontal dashed line at 0 represents no net change from baseline.

LYSC98 demonstrated potent efficacy against the *S. aureus* strains in our study. The maximal effect reached a 4.9 to 5.6 log_10_ CFU kill compared with the initial bacterial burden. The dose-response data were modeled using the sigmoid E_max_ equation, showing *f*C_max_/MIC was a strong predictor of treatment outcomes base on regression analysis (**Figure 6**, R^2^ 0.9585, R^2^ 0.8952, respectively).

**FIGURE 6.**
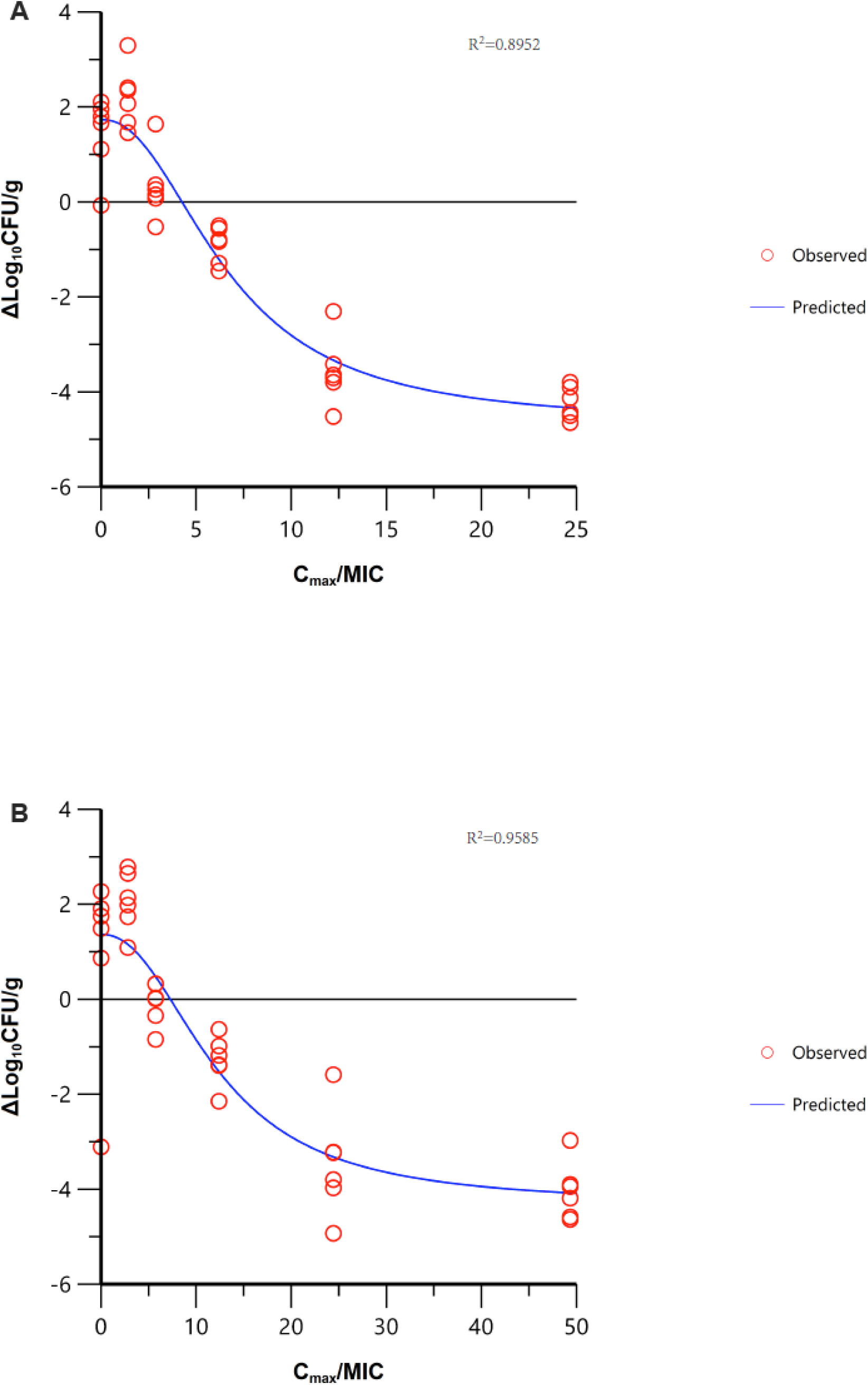
Correlation between PK/PD index fC_max_/MIC and efficacy of LYSC98 against two clinical *S. aureus* strains using a neutropenic murine thigh infection model in dose-escalation experiment. Treatment was initiated at 2 h post infection. LYSC98 was intravenous administered with single dose range of 1-16 mg/kg. Each point represents data for each sample. R^2^, square of the correlation coefficient.

The *f*C_max_/MIC values necessary to produce a target bacterial burden are shown in **Table 4**. Briefly, the median *f*C_max_/MIC targets needed for static, 1-log_10_CFU, 2-log_10_CFU, 3- log_10_CFU and 4-log_10_CFU were 5.78, 8.17, 11.14, 15.85, 30.58, respectively.

**TABEL 4.** In vivo activity and PK/PD analysis of LYSC98 against clinical organisms.

| Strains | $C_{\max}/MIC$ | | | | |
| --- | --- | --- | --- | --- | --- |
|  | Static | -1 log | -2 log | -3 log | -4 log |
| 18-W26-14 | 4.25 | 5.83 | 7.81 | 10.73 | 17.76 |
| 18-W27-73 | 7.31 | 10.51 | 14.46 | 20.97 | 43.39 |
| Mean | 5.78 | 8.17 | 11.14 | 15.85 | 30.58 |
| SD | 2.2 | 3.3 | 4.7 | 7.2 | 18.1 |

## DISCUSSION

An ideal optimization requires a good knowledge of the mechanisms involved in the effect of the antibiotics (pharmacodynamics, PD) and the change of the antibiotic concentration in the body of the patients (pharmacokinetics, PK). Pharmacokinetic/Pharmacodynamic (PK/PD) analysis integrates them and studies the dosing required to enhance the possibility of success of the antibiotic therapy, as well as minimize the side effects and the emergence of resistances (Asin-Prieto et al., 2015; Gabrielsson et al., 2010). Using a neutropenic murine thigh infection model, our study determined the magnitudes of *f*C_max_/MIC of LYSC98, a new antibacterial compound, associated with various levels of bacterial reduction for *S. aureus* strains. Since there is no available PK data from human, we did not choose Monte Carlo Simulation to evaluated the clinical dosing regimens or PK/PD breakpoints. This work should be further completed in the future development and research.

The inclusion of multiple bacterial isolates with various susceptibility should be considered in animal PK/PD experiment design to obtain a robust PK/PD target (Bulitta et al., 2019; Li et al., 2021). In our study, we used one ATCC strain and two clinical strains with MIC of 2-4mg/L to LYSC98. Only two clinical strains were used, which was the deficiency of this study. However, according to the existing in vitro MIC results, the two strains selected were STRAINS MIC_50_ and MIC_90_ of LYSC98, so the results were fairly representative. We also referred to the MIC range of these strains to vancomycin (0.5-1mg/L), and the MIC_90_ of vancomycin is 1mg according to previous studies (Liang et al., 2018). Therefore, the strains we choose are representative in *S. aureus* strains. Besides, MSSA and MRSA were included in our study which indicated the weak impact of penicillin resistance on LYSC98 activity. There was a difference in the MIC values of the strains while the bactericidal effect of LYSC98 were similar, it may be caused by the differences in virulence, tolerance and adaptive capacity to hosts between the strains. It also illustrates the importance to adopt a PK/PD approach combining in vivo and in vitro studies for dosing regimen evaluation rather than based on MIC values alone (Velkov et al., 2013).

The PK/PD indices of vancomycin antibiotics is generally considered as *f*AUC_0-24_MIC (Moise-Broder et al., 2004; Nielsen et al., 2011). Some other researchers think that there are no significant correlations between PK/PD indices and the clinical or microbiological efficacy of vancomycin (Shen et al., 2018). While in our study, E_max_ model analysis showed that for vancomycin derivate LYSC98, *f*C_max_/MIC (R^2^ 0.8941) correlated better with efficacy rather than *f*AUC_0-24_MIC (R^2^ 0.7793) and was only weakly correlated with %T>MIC (R^2^ 0.5424).

PK/PD indices are the best descriptors of antibiotic efficacy depending on the activity pattern of each antibiotic (Reed, 2000). As a glycopeptide antibiotic, the mechanism of vancomycin is generally believed to be that it binds to alanine at the end of the precursor of the sensitive bacterial cell wall and blocks the synthesis of peptidoglycan, thus leading to cell wall defects and killing bacteria(Wilhelm, 1991). In the common classification, vancomycin antibiotics belong to antibacterial agents with concentration independent killing and long-term persistence (Rybak, 2006). Due to the prolonged persistent effects that protect against regrowth when active drug concentration falls below the MIC, the best PK/PD indexes for these drugs are *f*C_max_/MIC or the *f*AUC_0-24_/MIC, which is closely consistent with our results (Asin-Prieto et al., 2015; Holmes, 2020).

As for the choices between *f*C_max_/MIC and *f*AUC_0-24_/MIC, we infer that the differences may be mainly considered from three aspects:

First, the half-life of different compounds needs to be considered. LYSC98 was engineered to have a significantly longer half-life in mice than vancomycin, which could explain its long-acting bactericidal effect. If the drug has a long half-life or postantibiotic effect (PAE), since %*f*T > MIC is easy to reach a high level, increasing the drug concentration could improve the effect, which mainly shows the concentration dependent characteristics. For instance, animal PK/PD studies of penicillin and amikacin, as well as some in vitro studies, showed an increased correlation of antibacterial effect with *f*C_max_/MIC and *f*AUC_0-24_/MIC when the half-lives of the drug were prolonged, and a shortened half-life is associated with %*f*T > MIC better(Craig et al., 1991; Nielsen et al., 2011).

Second, doses may also be an important factor. The same drug may have different PK/PD indices at different dose levels(Tam and Nikolaou, 2011). In our study, we used a total of 3 doses of 2-8mg/kg LYSC98 to conduct pharmacodynamic experiments. The data were well linear, but there was still a possibility that the maximum efficacy was not covered. This possibility may lead to a smaller overall dose of administration, a greater dependence of efficacy on concentration, and the PK/PD index will be more inclined to *f*C_max_/MIC. It may be a limitation of our study.

Third, we need to consider the impact of additional antimicrobial mechanisms. Based on previous study, vancomycin can also change the permeability of bacterial cell membrane, and selectively inhibit the synthesis of bacterial RNA(Okano et al., 2017). Ratio on different antibacterial mechanisms of LYSC98 may also have changed with specific structural modifications. One potent evidence is that it works well against vancomycin-resistant enterococcus (**Supplementary Table 1**). Certainly, this explanation remains hypothesis until further pharmacological confirmation.

## CONCLUSION

LYSC98, a novel vancomycin derivative, showed effective bactericidal efficacy against *Staphylococcus aureus* and reduced the bacteria loading in a murine thigh infection model at a dose of 2-8mg/kg. Different PK/PD indices were used to fit the pharmacodynamic parameters, and the results showed that *f*C_max_/MIC fitting results were the most appropriate indices, which could better represent LYSC98’s bactericidal properties. The *f*C_max_/MIC target values required to achieve static, −1 log, −2 log, −3 log and −4 log antibacterial activity in mice infected thigh were 5.78, 8.17, 11.14, 15.85 and 30.58, respectively. Our study provides effective preclinical data for further study and rational application of LYSC98.

## Supplementary Information

**Supplementary Table 1.** Minimum inhibitory concentration (MIC) of LYSC98 in different *S. aureus*

| NO. | Bacteria types | Vancomycin | Teicoplanin | LYSC98 |
| --- | --- | --- | --- | --- |
| 1 | WT | 0.5 | - | 1 |
| 2 | WT | 0.5 | - | 2 |
| 3 | MRSA | 0.5 | 0.5 | 2 |
| 4 | MRSA | 0.5 | 0.5 | 2 |
| 5 | MRSA | 0.5 | 0.5 | 2 |
| 6 | MRSA | 0.5 | 0.5 | 2 |
| 7 | MRSA | 0.5 | 0.5 | 2 |
| 8 | MRSA | 0.5 | 0.25 | 1 |
| 9 | MRSA | 0.5 | 0.5 | 2 |
| 10 | MRSA | 0.5 | 4 | 2 |
| 11 | MRSA | 0.5 | 0.5 | 2 |
| 12 | MRSA | 0.5 | 1 | 2 |
| 13 | MRSA | 0.5 | 1 | 2 |
| 14 | MRSA | 0.5 | 1 | 2 |
| 15 | MRSA | 0.5 | 1 | 2 |
| 16 | MRSA | 0.5 | 0.5 | 1 |
| 17 | MRSA | 0.5 | 1 | 2 |
| 18 | MRSA | 0.5 | 0.5 | 2 |
| 19 | MRSA | 0.5 | 1 | 2 |
| 20 | MRSA | 0.5 | 0.5 | 2 |
| 21 | MRSA | 0.5 | 0.5 | 2 |
| 22 | MRSA | 0.5 | 1 | 2 |
| 23 | MRSA | 0.5 | 0.25 | 2 |
| 24 | MRSA | 1 | 0.5 | 4 |
| 25 | MRSA | 1 | 0.25 | 2 |
| 26 | MRSA | 0.5 | 0.5 | 2 |
| 27 | MRSA | 0.5 | 0.125 | 1 |
| 28 | MRSA | 0.5 | 0.125 | 2 |
| 29 | MRSA | 0.5 | 0.5 | 1 |
| 30 | MRSA | 0.5 | 0.5 | 4 |
| 31 | VRSA | 16 | - | 2 |
| 32 | VRSA | 8 | - | 2 |
| 33 | VRSA | 16 | - | 2 |
| 34 | VRSA | 16 | - | 4 |

WT, wild type; MRSA, methicillin-resistant *Staphylococcus aureus*; VRSA, vancomycin-resistant *S. aureus*. The MIC unit is “mg/mL”.

**Supplementary Table 2.** In vivo protective effect of LYSC98 on MRSA infected mice (ED50)

| Groups | Drugs | Deli. | Dose<br>(mg/kg) | Conc.<br>(mg/ml) | N | Deaths | M.R.<br>(%) | ED<sub>50</sub><br>95% CI<br>(mg/kg) |
| --- | --- | --- | --- | --- | --- | --- | --- | --- |
| Treated<br>(3×10<sup>6</sup>CFU/ml) | Linezolid | i.v. | 10 | 0.4 | 8 | 1 | 12.5 | 4.64<br>3.07-7.60 |
|  |  |  | 5 | 0.2 | 8 | 4 | 50.0 |  |
|  |  |  | 2.5 | 0.1 | 8 | 6 | 75.0 |  |
|  |  |  | 1.25 | 0.05 | 8 | 8 | 100 |  |
|  | Vancomycin | i.v. | 10 | 0.4 | 8 | 1 | 12.5 | 3.58<br>2.32-5.84 |
|  |  |  | 5 | 0.2 | 8 | 2 | 25.0 |  |
|  |  |  | 2.5 | 0.1 | 8 | 6 | 75.0 |  |
|  |  |  | 1.25 | 0.05 | 8 | 7 | 87.5 |  |
|  |  |  | 0.6 | 0.025 | 8 | 8 | 100.0 |  |
|  | LYSC-98 | i.v. | 10 | 0.4 | 8 | 0 | 0 | 1.11<br>0.41-1.86 |
|  |  |  | 5 | 0.2 | 8 | 1 | 12.5 |  |
|  |  |  | 2.5 | 0.1 | 8 | 2 | 25.0 |  |
|  |  |  | 1.25 | 0.05 | 8 | 3 | 37.5 |  |
|  |  |  | 0.6 | 0.025 | 8 | 6 | 75.0 |  |
| Untreated<br>(3×10<sup>5</sup>CFU/ml) | - | - | - | - | 8 | 8 | 100.0 | - |
| Control | - | - | - | - | 8 | 0 | 0 | - |

M.R., mortality rate; i.v., intravenous injection.

## REFERENCES

Asempa, T.E., Abdelraouf, K., Carabeo, T., Schuch, R., and Nicolau, D.P. (2020). Synergistic Activity of Exebacase (CF-301) in Addition to Daptomycin against Staphylococcus aureus in a Neutropenic Murine Thigh Infection Model. Antimicrob Agents Chemother 64.

Asin-Prieto, E., Rodriguez-Gascon, A., and Isla, A. (2015). Applications of the pharmacokinetic/pharmacodynamic (PK/PD) analysis of antimicrobial agents. J Infect Chemother 21, 319–329.

Bulitta, J.B., Hope, W.W., Eakin, A.E., Guina, T., Tam, V.H., Louie, A., Drusano, G.L., and Hoover, J.L. (2019). Generating Robust and Informative Nonclinical In Vitro and In Vivo Bacterial Infection Model Efficacy Data To Support Translation to Humans. Antimicrob Agents Chemother 63.

Chambers, H.F. (1997). Methicillin resistance in staphylococci: molecular and biochemical basis and clinical implications. Clin Microbiol Rev 10, 781–791.

Craig, W.A., Redington, J., and Ebert, S.C. (1991). Pharmacodynamics of amikacin in vitro and in mouse thigh and lung infections. J Antimicrob Chemother 27 Suppl C, 29–40.

David, M.Z., and Daum, R.S. (2010). Community-associated methicillin-resistant Staphylococcus aureus: epidemiology and clinical consequences of an emerging epidemic. Clin Microbiol Rev 23, 616–687.

Drebes, J., Kunz, M., Pereira, C.A., Betzel, C., and Wrenger, C. (2014). MRSA infections: from classical treatment to suicide drugs. Curr Med Chem 21, 1809–1819.

Gabrielsson, J., Green, A.R., and Van der Graaf, P.H. (2010). Optimising in vivo pharmacology studies--Practical PKPD considerations. J Pharmacol Toxicol Methods 61, 146–156.

Growcott, E.J., Cariaga, T.A., Morris, L., Zang, X., Lopez, S., Ansaldi, D.A., Gold, J., Gamboa, L., Roth, T., Simmons, R.L., et al. (2019). Pharmacokinetics and pharmacodynamics of the novel monobactam LYS228 in a neutropenic murine thigh model of infection. J Antimicrob Chemother 74, 108–116.

Hidayat, L.K., Hsu, D.I., Quist, R., Shriner, K.A., and Wong-Beringer, A. (2006). High-dose vancomycin therapy for methicillin-resistant Staphylococcus aureus infections: efficacy and toxicity. Arch Intern Med 166, 2138–2144.

Holmes, N.E. (2020). Using AUC/MIC to guide vancomycin dosing: ready for prime time? Clin Microbiol Infect 26, 406–408.

Howe, R.A., Bowker, K.E., Walsh, T.R., Feest, T.G., and MacGowan, A.P. (1998). Vancomycin-resistant Staphylococcus aureus. Lancet 351, 602.

Lakhundi, S., and Zhang, K. (2018). Methicillin-Resistant Staphylococcus aureus: Molecular Characterization, Evolution, and Epidemiology. Clin Microbiol Rev 31.

Lepak, A.J., Zhao, M., Marchillo, K., VanHecker, J., and Andes, D.R. (2019). In Vivo Pharmacodynamics of Omadacycline against Staphylococcus aureus in the Neutropenic Murine Thigh Infection Model. Antimicrob Agents Chemother 63.

Li, X., Chen, Y., Xu, X., Li, Y., Fan, Y., Liu, X., Bian, X., Wu, H., Zhao, X., Feng, M., et al. (2021). Pharmacokinetics and Pharmacodynamics of Nemonoxacin in a Neutropenic Murine Lung Infection Model Against Streptococcus Pneumoniae. Front Pharmacol 12, 658558.

Liang, X., Fan, Y., Yang, M., Zhang, J., Wu, J., Yu, J., Tao, J., Lu, G., Zhang, H., Wang, R., et al. (2018). A Prospective Multicenter Clinical Observational Study on Vancomycin Efficiency and Safety With Therapeutic Drug Monitoring. Clin Infect Dis 67, S249–S255.

Lowy, F.D. (1998). Staphylococcus aureus infections. N Engl J Med 339, 520–532.

McGuinness, W.A., Malachowa, N., and DeLeo, F.R. (2017). Vancomycin Resistance in Staphylococcus aureus. Yale J Biol Med 90, 269–281.

Michel, M., and Gutmann, L. (1997). Methicillin-resistant Staphylococcus aureus and vancomycin-resistant enterococci: therapeutic realities and possibilities. Lancet 349, 1901–1906.

Moise-Broder, P.A., Forrest, A., Birmingham, M.C., and Schentag, J.J. (2004). Pharmacodynamics of vancomycin and other antimicrobials in patients with Staphylococcus aureus lower respiratory tract infections. Clin Pharmacokinet 43, 925–942.

Mouton, J.W., Brown, D.F., Apfalter, P., Canton, R., Giske, C.G., Ivanova, M., MacGowan, A.P., Rodloff, A., Soussy, C.J., Steinbakk, M., et al. (2012). The role of pharmacokinetics/pharmacodynamics in setting clinical MIC breakpoints: the EUCAST approach. Clin Microbiol Infect 18, E37–45.

Nielsen, E.I., Cars, O., and Friberg, L.E. (2011). Pharmacokinetic/pharmacodynamic (PK/PD) indices of antibiotics predicted by a semimechanistic PKPD model: a step toward model-based dose optimization. Antimicrob Agents Chemother 55, 4619–4630.

Okano, A., Isley, N.A., and Boger, D.L. (2017). Total Syntheses of Vancomycin-Related Glycopeptide Antibiotics and Key Analogues. Chem Rev 117, 11952–11993.

Rao, Q., Shang, W., Hu, X., and Rao, X. (2015). Staphylococcus aureus ST121: a globally disseminated hypervirulent clone. J Med Microbiol 64, 1462–1473.

Reed, M.D. (2000). Optimal antibiotic dosing. The pharmacokinetic-pharmacodynamic interface. Postgrad Med 108, 17–24.

Rybak, M.J. (2006). The pharmacokinetic and pharmacodynamic properties of vancomycin. Clin Infect Dis 42 Suppl 1, S35–39.

Shen, K., Yang, M., Fan, Y., Liang, X., Chen, Y., Wu, J., Yu, J., Zhang, H., Wang, R., Zhang, F., et al. (2018). Model-based Evaluation of the Clinical and Microbiological Efficacy of Vancomycin: A Prospective Study of Chinese Adult In-house Patients. Clin Infect Dis 67, S256–S262.

Tam, V.H., and Nikolaou, M. (2011). A novel approach to pharmacodynamic assessment of antimicrobial agents: new insights to dosing regimen design. PLoS Comput Biol 7, e1001043.

Velkov, T., Bergen, P.J., Lora-Tamayo, J., Landersdorfer, C.B., and Li, J. (2013). PK/PD models in antibacterial development. Curr Opin Microbiol 16, 573–579.

Wilhelm, M.P. (1991). Vancomycin. Mayo Clin Proc 66, 1165–1170.

Zhou, K., Li, C., Chen, D., Pan, Y., Tao, Y., Qu, W., Liu, Z., Wang, X., and Xie, S. (2018). A review on nanosystems as an effective approach against infections of Staphylococcus aureus. Int J Nanomedicine 13, 7333–7347.

